# Mutational insights into human kynurenine aminotransferase 1: modulation of transamination and β-elimination activities across diverse substrates

**DOI:** 10.1101/2025.03.27.645645

**Authors:** Arun Kumar Selvam, Renhua Sun, Ali Razaghi, Hugh Salter, Tatiana Sandalova, Mikael Björnstedt, Adnane Achour

## Abstract

Human kynurenine aminotransferase 1 (hKYAT1) plays a crucial role in the transamination of aromatic amino acids and kynurenine. This promiscuous homodimeric enzyme transaminates various amino acids into their corresponding alpha-keto acids. Additionally, hKYAT1 is known to catalyze the beta-elimination of cysteine-S-conjugates and cysteine-Se-conjugates. In this study, we performed mutational analyses of hKYAT1, targeting its catalytic, ligand-binding, and substrate-binding sites. The transamination activity of thirteen mutant variants was systematically evaluated against sixteen different amino acid substrates, including kynurenine, selenomethionine (SeMet), and Se-methylselenocysteine (MSC), as well as for the β-elimination of SeMet and MSC. Our results demonstrate that mutations of residues E27 in the catalytic site and H279 in the substrate-stabilizing site significantly enhanced the transamination of several amino acids, including phenylalanine, tryptophan, histidine, and MSC. The H279F mutation increased transamination and β-elimination of MSC by two- and 1.5-fold, respectively. Furthermore, mutation at the ligand-binding residues R398 and N185 abolished the transamination activity of hKYAT1. Interestingly, none of the tested mutations affected the transamination of L-kynurenine, a natural substrate of hKYAT1. These findings provide a foundation for the rational design of selective inhibitors with potential therapeutic applications.

## Introduction

The metabolism of amino acids is essential for the proliferation and maintenance of all living cells. Several amino acids serve as neurotransmitters, regulate the cellular redox status, and are involved in critical pathways including the activation of the mTOR pathway or DNA methylation (1, 2). Notably, rapidly proliferating cells, *i*.*e*. immune or tumor cells, exhibit a greater demand for amino acids compared to normal cells. In addition, the activation of aerobic glycolysis, a key metabolic reprogramming process in cancer, increases reliance on glutamine as an energy source (1, 2). Therefore, the anabolic and catabolic regulation of amino acids is essential for normal cellular function and constitutes a critical pathway in numerous diseases.

Transamination is a pivotal step in the catabolism of a large ensemble of amino acids. To date, at least six distinct human cytosolic transaminases have been identified, all involved in amino acid catabolic pathways (3). Notably, cytosolic kynurenine aminotransferase (KYAT1) catalyzes the conversion of kynurenine to kynurenic acid (KYNA), a key reaction within the kynurenine pathway (KP), the primary catabolic route for tryptophan. The KP comprises a complex cascade of reactions catalyzed by eight different enzymes (4). Tryptophan metabolism occurs in most cells and plays important roles in various organs, including the kidneys, liver, leucocytes, heart, lungs, and cells in the brain such as astrocytes and microglia (5).

The KYAT1 enzyme is part of the type I aminotransferase subfamily, with aspartate aminotransferase as its prototypical member (6), commonly used as a liver damage marker (7). The activity of aspartate aminotransferases is dependent on the cofactor pyridoxal 5’-phosphate (PLP), which enables the transfer of the amine group from the substrate to the formed α-keto-acid (8). To date, four human KYAT variants (hKYAT1-hKYAT4) have been identified (4), with broad substrate specificity. Additionally, KYAT1 and KYAT3 exhibit β-lyase activity toward cysteine-S-conjugate and cysteine-Se-conjugate substrates that possess strong electron-withdrawing groups at the sulfur or selenium atom (9).

Notably, *in vitro* studies have demonstrated that hKYAT1 exhibits broad substrate specificity (9), achieving its highest catalytic efficiency with L-glutamine, L-phenylalanine, L-leucine, L-tryptophane, L-kynurenine, and L-methionine (10). Additionally, it also exhibits significant β-lyase activity with S-(1,2-dichlorovinyl)-L-cysteine (DCVC), S-(1,1,2,2-tetrafluoroethyl)-L-cysteine (TFEC) and MSC (11). This broad functional capacity accounts for the various names attributed to hKYAT1, such as kynurenine aminotransferase 1 (EC 2.6.1.7), glutamine transaminase K (GTK) (EC 2.6.1.64), and cysteine conjugate β-lyase 1 (EC 4.4.1.13).

hKYAT1 has been implicated in various diseases, including cancer (12), neurodegenerative and psychiatric disorders (13), inflammation, and obesity (14), primarily due to its role in tryptophan metabolism within the KP. Altered hKYAT1 activity within the KP has been associated with conditions such as cerebral ischemia (15), while significantly reduced hKYAT1 mRNA expression has been linked to severe skeletal disorders (16). Notably, large amounts of compounds produced along the KP play important roles in modulating different biological processes within the organism (17-20). However, the significance of KYAT1 extends beyond the KP, as it is a key component of the glutaminase II pathway. This enzyme catalyzes the transamination of glutamine to α-ketoglutaramate (KGM) in the presence of an α-keto acid acceptor. This reaction is coupled to ω-amidase, which hydrolyzes KGM to α-ketoglutarate, facilitating its entry into the tricarboxylic acid (TCA) cycle (11). Many cancers rely on increased glutamine catabolism for energy production, often due to the decoupling of the glycolytic pathway from the TCA cycle (21). Interestingly, both human KYAT1 and KYAT3 have been found to be overexpressed in various cancer cell lines (11, 22). Several crystal structures of the hKYAT1 holoenzyme and hKYAT1 in complex with an array of substrates or inhibitors have been previously determined (23-25). While more than 500 different point mutations have been identified in hKYAT1, none have been produced or functionally characterized. To date, no mutational studies have clarified the role of critical residues in hKYAT1. To our knowledge, the only reported variant is the naturally occurring E61G mutation in rat KYAT1 (rKYAT1), which leads to reduced aminotransferase activity and spontaneous hypertension in rats (26). It should be noted that residue E61 in rKYAT1 corresponds to E27 in hKYAT1. In this study, we present the first comprehensive mutational analysis of the substrate-binding region of hKYAT1. By evaluating the impact of various mutations on both transamination and β-elimination activities using a diverse range of substrates, we aimed to elucidate the molecular determinants involved in substrate recognition by hKYAT1. These findings provide a foundation for the rational design of selective inhibitors with potential therapeutic applications.

## Results

### Structure-based selection and design of mutations

Several crystal structures of hKYAT1 have been determined in complex with the co-factors PLP (PDB codes 1W7L (25), 3FVS (23) and 4WLH (24)), or pyridoxamine 5’-phosphate (PMP, 1W7N (25)), as well as substrates such as L-Phe (1W7M (25)), and inhibitors like indole-3-acetic acid (3FVU (23)). hKYAT1 forms a stable homodimer with a buried interface surface area of approximately 3000Å^2^ (**Figure 1A**). The substrate binds above the PLP ring, which is covalently linked to the lysine residue K247 in the active site of hKYAT1. This pocket is composed of residues from both subunits **(Figure 1B)**. In this study, we aimed to introduce subtle modifications that would preserve the enzyme’s function and structural integrity, while selectively altering active site residues. Our goal was to evaluate the impact of these substitutions on substrate specificity and enzymatic activity. As PLP is essential for hKYAT1 catalysis, we avoided mutating any residue involved in PLP binding and focused on those that contribute to substrate interactions.

**Figure 1.**
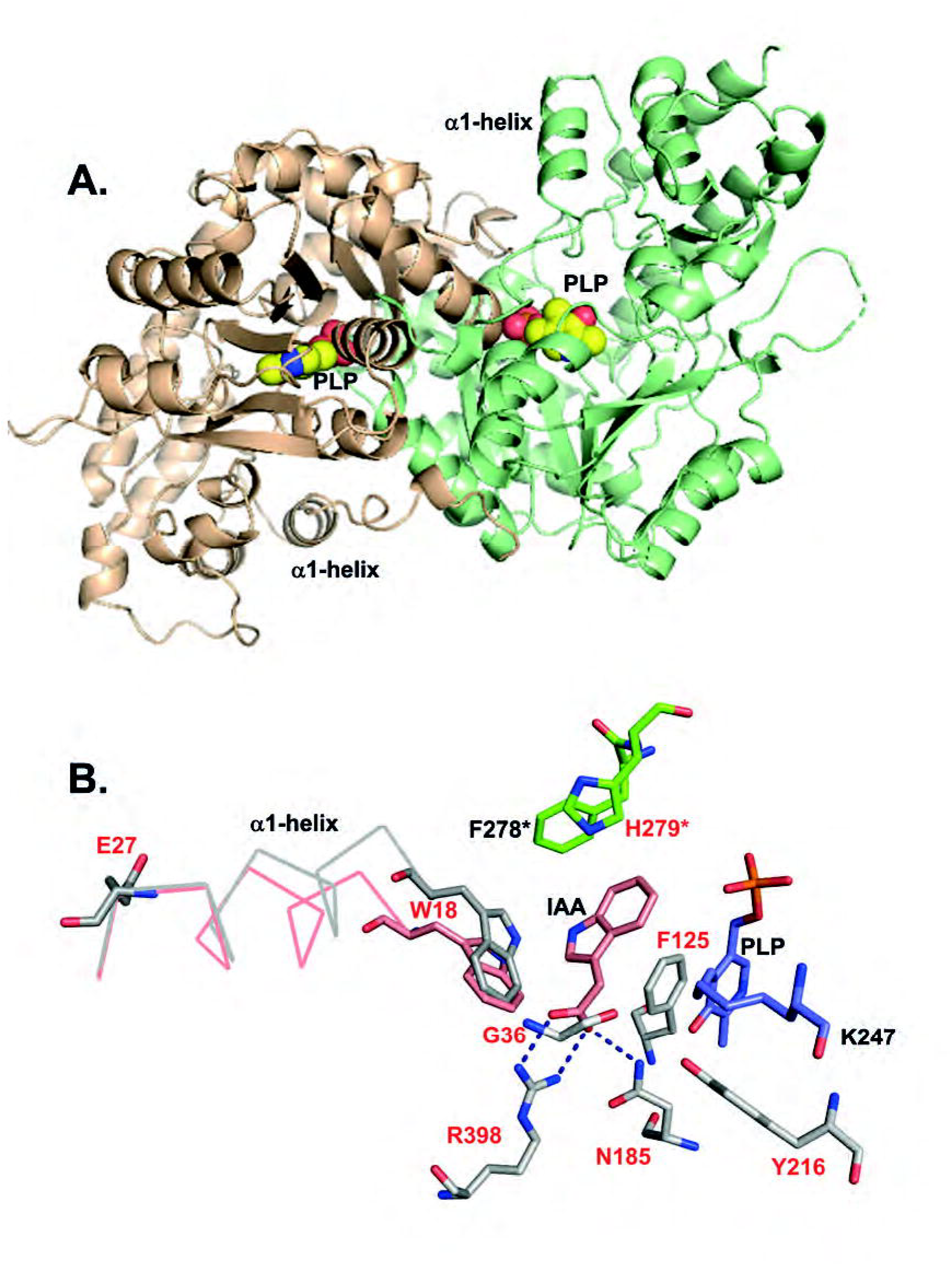
Structural basis for residue selection within the active site of hKYAT1. Crystal structures of hKYAT1 (PDB code 3FVS) (23) reveal that both subunits contribute to forming the enzyme’s active site. **A**. The two subunits are in pink and light green, with the active sites marked by the co-factor PLP. The N-terminal α1-helix (residues 18-27), crucial for the catabolic activity of hKYAT1 is also highlighted. **B**. Comparison of the crystal structures of ligand-free hKYAT1 (PDB code 3FVS, gray) and hKYAT1 in complex with the inhibitor IAC (PDB code 3FVS, pink) reveals a lateral shift of the α1-helix to accommodate ligand binding. Residues from the two hKYAT1 subunits are shown in gray and green. All residues mutated in this study are labeled in red.

The crystal structure of hKYAT1 complexed with L-Phe (25) or the inhibitor IAC (23) reveals the substrate positioned between residues F125 and H279* (**Figure 1B**, * indicates residues from the other subunit). Residues N185 and R398 form hydrogen bonds and salt bridges with the substrate’s α-carboxylate group, positioning its α-amino group above the PLP in a conformation conducive to catalysis. The ligand-binding site of hKYAT1 is composed of several aromatic residues, including W18, Y101, and F125 from one monomer, and Y63*, H279*, and F278* from the other subunit. To investigate the role of these aromatic residues, we introduced mutations in several key positions. We hypothesized that the F125H or H279F mutations would not prevent substrate binding but could alter or modulate the enzyme’s functional activity toward both aromatic and non-aromatic substrates. Additionally, the R398A mutation was introduced to determine whether the N185 residue alone could maintain the substrate in the correct position. The double mutation N185G/R398K was also introduced to evaluate whether lysine at position 398 could stabilize the substrate in the absence of support from N185. Furthermore, we replaced N185 with glutamine (N185Q) to reduce the size of the substrate-binding pocket without altering the side chain’s properties. Previous studies have shown that the tyrosine in aminotransferases that is analogous to Y216 in hKYAT1 can substitute for arginine 398 to anchor the substrate (23). To test this, we introduced the Y216R mutation, both individually and in combination with the R398A mutation (Y216R/R398A). Additionally, we examined the role of residue G36, located near the substrate-binding site. The G36S mutation aimed to decrease the size of the substrate-binding pocket and enhance substrate selectivity.

Unlike subgroup I aminotransferases, such as aspartate aminotransferase, which undergo conformational changes upon ligand binding to close the active site (8), hKYAT1 binds substrates without significant domain movement. Instead, substrate-binding involves a shift in the N-terminal α1-helix (residues P17-E27) with a pivotal movement facilitated by the glutamate residue E27. This shift tilts the α1-helix toward the active site and is accompanied by a rotation of Y101 (**Figure 1B**). The proximity between W18 and the substrate is reduced to 3.9 Å, creating optimal van der Waals interactions (**Figure 1B**). To assess the functional significance of the α1-helix movement, we introduced mutations at W18 and E27. We hypothesized that the E27G mutation would increase the lateral flexibility of the α1-helix, while replacing the bulky tryptophan residue W18 with smaller side chains (W18H, W18M, or W18L) would modify substrate specificity (**Figure 1B**).

All mutated residues examined in this study are listed in **Table S1**. Most experiments were conducted using cell lysates. To account for differences in expression levels among the mutants, Western blot analyses were performed. The majority of the mutants displayed expression levels similar to that of the wild-type except for H279F-, R398A-, N185Q-, F125H-, and W18H-hKYAT1 which exhibited two-fold higher expression (**Figure S1**). Notably, the W18L, W18M, and W18H mutants displayed varying expression levels, with W18H-hKYAT1 displaying the highest expression and W18L-hKYAT1 the lowest.

### Mutation-driven modulation of hKYAT1 activity: Divergent effects on aromatic amino acids and kynurenine

Residue substitutions in the catalytic site (helix α1, E27G), ligand binding site (N185Q), and substrate stabilization site (H279F) resulted in a 1.5-to two-fold increase in the transamination of L-Phe as compared to wild-type hKYAT1 lysate (**Figure 2A**). In contrast, other mutations significantly reduced L-Phe transamination (**Figure 2A**). The H279F substitution significantly increased the transamination activity of the recombinant hKYAT1, corroborating the results from cell lysate assays (**Figure 2B**). Interestingly, recombinant N185Q-hKYAT1 displayed significantly reduced transamination activity (**Figure 2B**), differing from the increased activity observed in cell lysates (**Figure 2A**). Substitutions at residue 398 (R398A- and N185G/R398K) abolished the transamination capacity of hKYAT1 for L-Phe (**Figure 2B**), as further supported by enzyme kinetics showing H279F with a higher K_cat_/K_m_ ratio compared to wild-type (**Figure 2C**). Aside from the observed dichotomy for the N185Q mutation, results from both cell lysate and recombinant protein assays were consistent. Additionally, the K_m_ and K_cat_ values for the five mutated enzyme variants support these findings (**Figure 2C, Table 1**) In conclusion, while some mutations significantly enhance transaminase activity, others completely abolish it, underscoring the critical roles of the selected residues in L-Phe transamination.

**Table 1:**
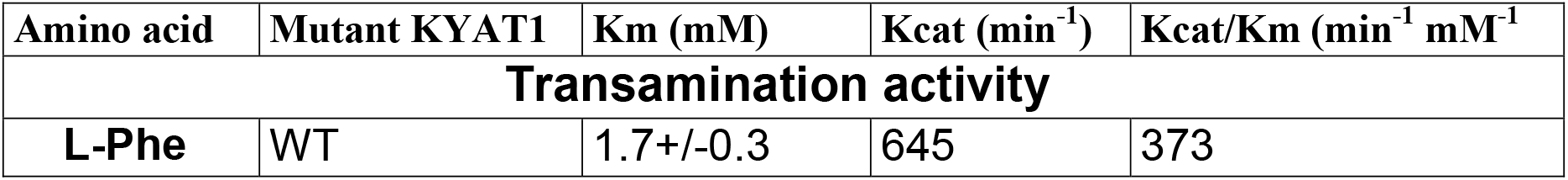
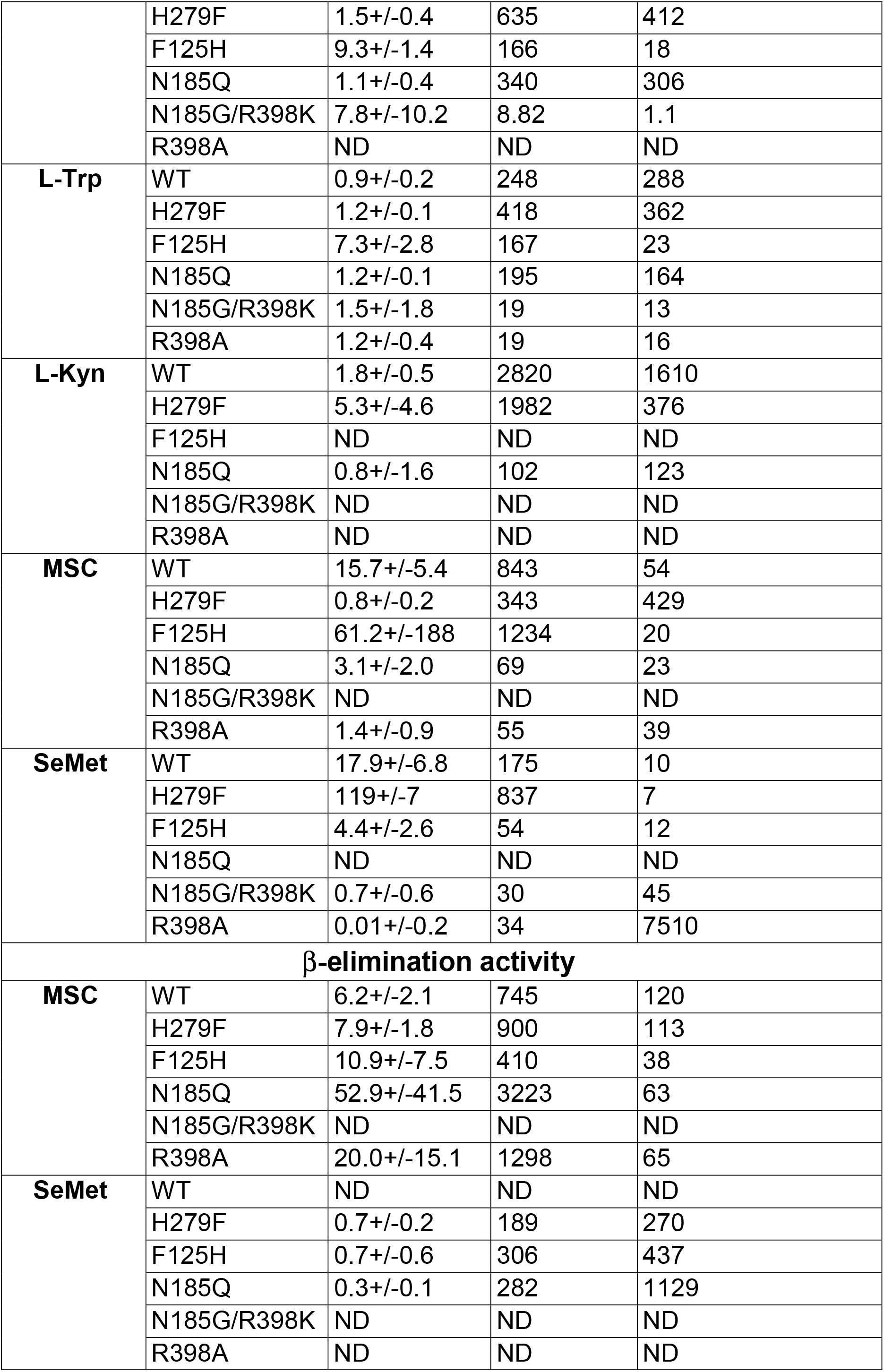
Michaelis-Menten kinetic parameters for transamination and β-elimination of the different recombinant mutant proteins with various substrates.

| Amino acid | Mutant KYAT1 | K<sub>m</sub> (mM) | K<sub>cat</sub> (min<sup>-1</sup>) | K<sub>cat</sub>/K<sub>m</sub> (min<sup>-1</sup> mM<sup>-1</sup>) |
| --- | --- | --- | --- | --- |
| **Transamination activity** |  |  |  |  |
| **L-Phe** | WT | 1.7+/-0.3 | 645 | 373 |
|  | H279F | 1.5+/-0.4 | 635 | 412 |
|  | F125H | 9.3+/-1.4 | 166 | 18 |
|  | N185Q | 1.1+/-0.4 | 340 | 306 |
|  | N185G/R398K | 7.8+/-10.2 | 8.82 | 1.1 |
|  | R398A | ND | ND | ND |
| **L-Trp** | WT | 0.9+/-0.2 | 248 | 288 |
|  | H279F | 1.2+/-0.1 | 418 | 362 |
|  | F125H | 7.3+/-2.8 | 167 | 23 |
|  | N185Q | 1.2+/-0.1 | 195 | 164 |
|  | N185G/R398K | 1.5+/-1.8 | 19 | 13 |
|  | R398A | 1.2+/-0.4 | 19 | 16 |
| **L-Kyn** | WT | 1.8+/-0.5 | 2820 | 1610 |
|  | H279F | 5.3+/-4.6 | 1982 | 376 |
|  | F125H | ND | ND | ND |
|  | N185Q | 0.8+/-1.6 | 102 | 123 |
|  | N185G/R398K | ND | ND | ND |
|  | R398A | ND | ND | ND |
| **MSC** | WT | 15.7+/-5.4 | 843 | 54 |
|  | H279F | 0.8+/-0.2 | 343 | 429 |
|  | F125H | 61.2+/-188 | 1234 | 20 |
|  | N185Q | 3.1+/-2.0 | 69 | 23 |
|  | N185G/R398K | ND | ND | ND |
|  | R398A | 1.4+/-0.9 | 55 | 39 |
| **SeMet** | WT | 17.9+/-6.8 | 175 | 10 |
|  | H279F | 119+/-7 | 837 | 7 |
|  | F125H | 4.4+/-2.6 | 54 | 12 |
|  | N185Q | ND | ND | ND |
|  | N185G/R398K | 0.7+/-0.6 | 30 | 45 |
|  | R398A | 0.01+/-0.2 | 34 | 7510 |
| **β-elimination activity** |  |  |  |  |
| **MSC** | WT | 6.2+/-2.1 | 745 | 120 |
|  | H279F | 7.9+/-1.8 | 900 | 113 |
|  | F125H | 10.9+/-7.5 | 410 | 38 |
|  | N185Q | 52.9+/-41.5 | 3223 | 63 |
|  | N185G/R398K | ND | ND | ND |
|  | R398A | 20.0+/-15.1 | 1298 | 65 |
| **SeMet** | WT | ND | ND | ND |
|  | H279F | 0.7+/-0.2 | 189 | 270 |
|  | F125H | 0.7+/-0.6 | 306 | 437 |
|  | N185Q | 0.3+/-0.1 | 282 | 1129 |
|  | N185G/R398K | ND | ND | ND |
|  | R398A | ND | ND | ND |

**Figure 2.**
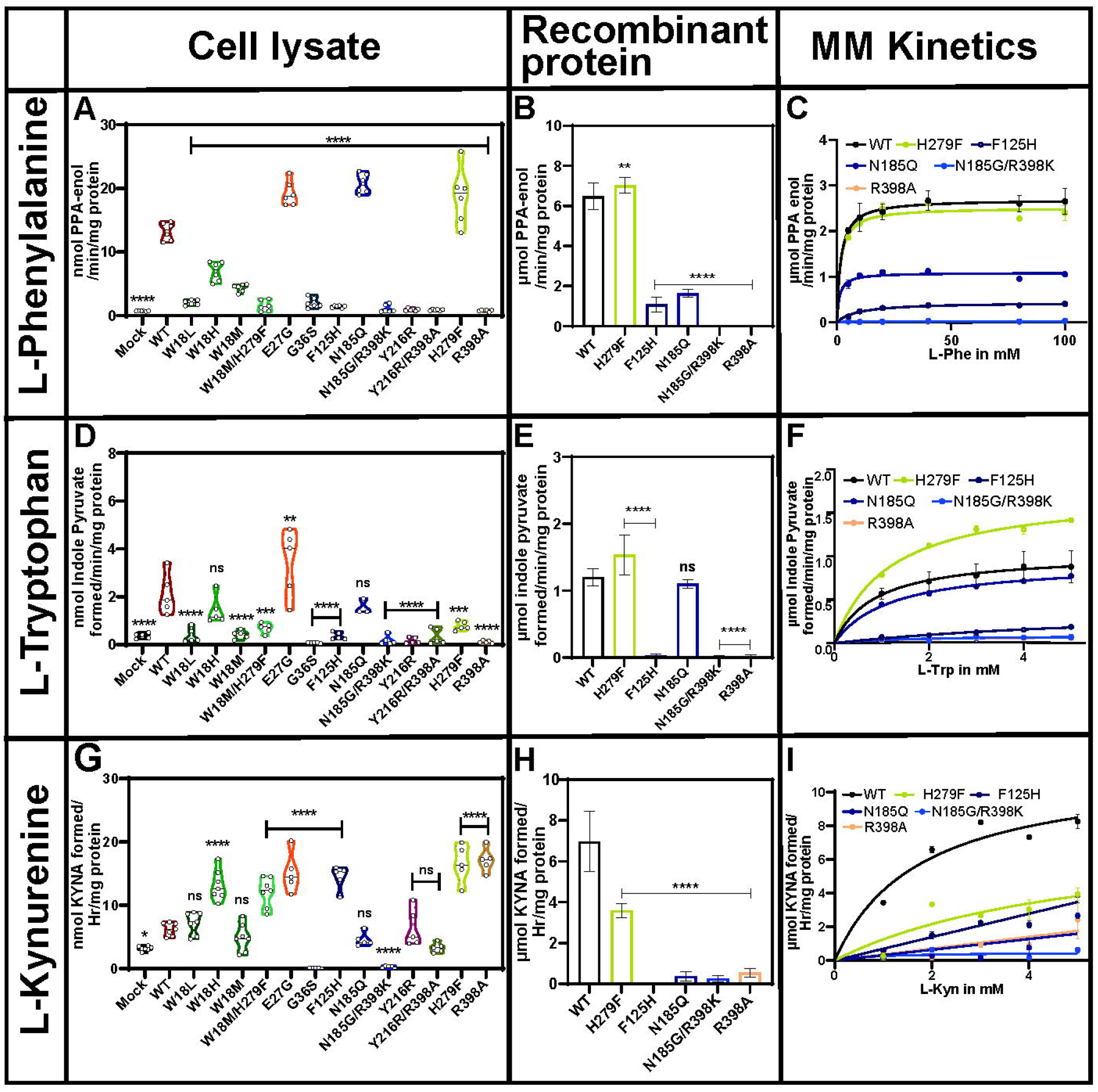
H279 mutation in the substrate stabilization site alters hKYAT1 transamination activity for aromatic amino acids and kynurenine. The transamination efficacy of thirteen hKYAT1 mutants was assessed for L-Phe (20mM), L-Trp (3mM), and Kyn (3mM) using whole-cell lysates with 20 µg of crude protein (**A, E, I**). Five recombinant hKYAT1 mutants were analyzed separately (**B, F, J**). The Michaelis-Menten kinetics for the wild-type and these five mutants are shown in **C, G**, and **K** for L-Phe, L-Trp, and L-Kyn, respectively. Statistical significance was determined using one-way ANOVA with a 95% confidence interval, followed by Dunnett’s multiple comparison test (mean ± sd *p<0.05, n ≥5 per group).

Tryptophan is another important aromatic amino acid substrate for hKYAT1. Most of the tested mutants either abolished or reduced hKYAT1’s transamination activity towards L-Trp, except for the E27G mutation, which resulted in a two-fold increase in activity compared to wild-type hKYAT1 lysate (**Figure 2D**).

Notably, the W18H and N185Q mutations did not affect L-Trp transamination in cell lysates as compared to the wild-type (**Figure 2D**). As observed for L-Phe, the results from recombinant proteins both confirmed and contrasted with the cell lysate data (**Figures 2D, 2E**). While the H279F variant displayed a 1.5-fold increase in activity compared to wild-type hKYAT1 in recombinant protein assays, no significant difference was observed in the cell lysates (**Figures 2D, 2E**). Interestingly, the recombinant protein results showed similar mutational effects for both L-Phe and L-Trp (**Figures 2B, 2E**), aligning well with the K_m_ and K_cat_ values for these substrates (**Figures 2C, 2F, Table 1**). The same set of mutations was tested with dL-Tyr (**Figures S2A, S2B**). The E27G mutation exhibited comparable transamination activity to wild-type hKYAT1 for dL-Tyr, while all other mutations reduced hKYAT1’s activity towards L-Tyr. These findings were confirmed by recombinant assays, where most mutations abolished dL-Tyr transamination, except for H279F (**Figures S2A, S2B**).

A major function of hKYAT1 is the transamination of L-kynurenine (Kyn) to kynurenic acid (Kyna). Interestingly, mutations W18H, W18M/H279F, E27G, F125H, H279F, and R398A increased the enzymatic activity of hKYAT1 toward L-Kyn by two-to four-fold in cell lysates (**Figure 2G**), a pattern that contrasts with the activity observed for L-Phe and L-Trp (**Figures 2A, 2D, and 2G**). However, all five tested recombinant mutants either reduced or abolished L-Kyna production as compared to the wild-type (**Figure 2H**). The K_m_ and K_cat_ values for the mutant recombinant hKYAT1 variants towards L-Kyn also differed significantly from those for L-Phe and L-Trp. Specifically, all five recombinant mutants, including H279F exhibited lower K_cat_/K_m_ ratios as compared to the wild-type, indicating lower catalytic efficiency for L-Kyn transamination (**Figures 2C, 2F, and 2I, Table 1**).

Overall, our results indicate that hKYAT1 employs different key residues for transamination depending on the substrate, whether it be aromatic amino acids or kynurenine. Despite some discrepancies between the results from cell lysates and recombinant proteins, we conclusively demonstrate that modifications at residue E27 lead to increased transamination activity across all aromatic substrates, including L-Kyn. Interestingly, the expression levels of the recombinant proteins did not directly correlate with hKYAT1 enzymatic activity (**Figure S1**). For instance, while H279F-hKYAT1 exhibited significantly higher transamination activity towards L-Phe, the F125H- and R398A-hKYAT1 variants showed no transamination capacity for these substrates. Even though the expression levels of F125H- and R398A-hKYAT1 were twice as high as wild-type-hKYAT1, they displayed no transamination activity across a broad range of tested substrates.

### Mutational analysis and substrate preferences of hKYAT1 highlight key residues in transamination activity

Amidic amino acids are essential for cancer cell survival as they can be converted into their respective α-keto acids via transamination. Notably, essential α-keto acids, such as α-ketoglutarate and oxaloacetate, which are derived from the transamination of glutamine and asparagine, respectively, play crucial roles in cell survival as key components of the tricarboxylic acid (TCA) cycle. Overexpression of hKYAT1 in HEPG2 cells resulted in a five-fold increase in L-Gln transamination compared to control cell lysates (**Figure 3**). However, mutations of residues in the ligand recognition site (W18), stabilization site (F125, Y216, and R398), and substrate stabilization site (H279) significantly reduced or completely abrogated enzymatic activity toward L-Gln. In contrast, mutations at the catalytic site (E27) increased hKYAT1 transamination activity toward L-Gln, while mutation of N185 in the ligand binding site showed activity as comparable to wild-type hKYAT1 (**Figure 3A**). These findings were consistent with the results obtained using recombinant proteins (**Figure 3B**). Furthermore, transamination activity toward L-Asn was minimal across all tested mutants using both cell lysates and recombinant proteins. However, mutations W18L and N185Q exhibited a notable increase in transamination activity as compared to wild-type hKYAT1 cell lysates (**Figure 3C**), and interestingly, recombinant F125H-hKYAT1 displayed a significant increase in transamination activity compared to wild-type hKYAT1 (**Figure 3D**). Similarly, hKYAT1 exhibited minimal activity with the acidic amino acid L-Asp, a finding consistent across both cell lysate and recombinant protein assays (**Figures S2C, S2D**). While some mutations suggested a modest increase in transamination activity in cell lysates, recombinant protein assays confirmed that L-Asn and L-Asp are not major substrates to hKYAT1 (**Figure 3D, Figure S2D**). Most hKYAT1 mutants displayed reduced transamination activity toward L-His in cell lysate assays, except for the E27G mutant, which showed enhanced activity (**Figure 3E**). Conversely, recombinant H279F displayed a significant increase in L-His transamination compared to wild-type hKYAT1, while all other mutations abrogated transamination activity (**Figure 3F**). These results underscore the importance of residue H279 in L-His transamination (**Figures 3E, 3F**).

**Figure 3.**
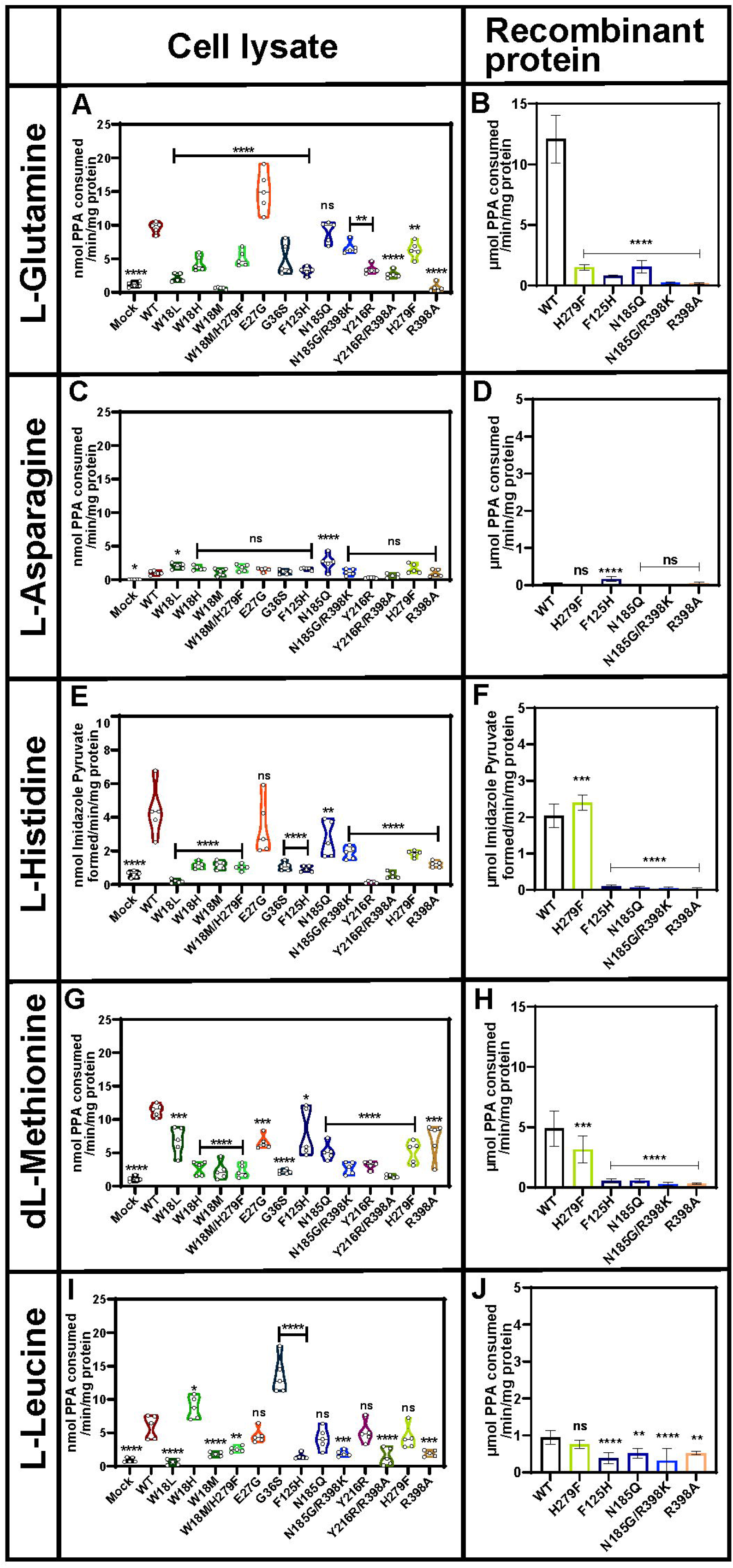
Mutations in hKYAT1 significantly reduce transamination activity with Gln and Met. Transamination efficacy of thirteen hKYAT1 mutant variants was tested for amidic L-Gln (3 mM), basic L-His (5 mM), acidic L-Asp (3 mM), sulfur-containing dL-Met (3 mM), and aliphatic L-Leu (3 mM). Panels **A, C, E, G** and **I** show results from cell lysates containing 20 µg of crude protein, while panels **B, D, F, H** and **J** present data from wild-type and five recombinant mutant proteins with 200 ng of recombinant protein. Statistical significance was determined with one-way ANOVA with a 95% confidence interval, followed by Dunnett’s multiple comparison test, (mean ± s.d., *P<0.05, n ≥5 per group).

Wild-type hKYAT1 overexpression had a profound impact on the transamination of the sulfur-containing amino acid methionine, resulting in a nine-fold increase in transamination activity as compared to control lysates (**Figure 3G**). Interestingly, all tested mutants significantly reduced hKYAT1’s transamination activity towards dL-Met in cell lysates (**Figure 3G**). This was further confirmed in recombinant protein assays, where all mutations, except H279F, completely abrogated dL-Met transamination (**Figure 3H**). Additionally, hKYAT1 displayed minimal activity with L-Cyss, a result corroborated by both cell lysate and recombinant protein assays (**Figures S2E, S2F**). hKYAT1 efficiently transaminated the aliphatic amino acid L-Leu, exhibiting a seven-fold increase in transamination activity as compared to control lysates (**Figure 3I**). Notably, the W18H and G36S mutations significantly increased transamination activity towards L-Leu compared to wild-type hKYAT1 (**Figure 3I**). In contrast, mutation of residue W18 to leucine (W18L) or methionine (W18M) reduced transamination activity toward L-Leu (**Figure 3I**). Several other mutations, including R398A, N185G/R398K, and Y216R/R398A, also reduced or abrogated L-Leu transamination, highlighting the importance of R398 in L-Leu transamination (**Figure 3I**). These results were validated by recombinant protein assays (**Figure 3J**). Additionally, we tested smaller hydrophobic substrates such as L-Ala, Pro, and Gly. While some transamination activity was detected, these residues are not primary substrates for hKYAT1 (**Figures S2G, S2H, S2I, S2J, S2K, S2L**). In conclusion, our results demonstrate that Gln, Met, His, and Leu are the most favorable substrates for hKYAT1, while Asn, Asp, Cyss, Ala, Pro, and Gly are the least preferred substrates.

### Functional characterization of hKYAT1 mutants: Enhanced activity with MSC compared to SeMet

KYAT1 transaminates Se-conjugated compounds, including MSC and SeMet, converting them to β-methylselenopyruvate (MSP) and α-keto-γ-methylselenobutyrate (KMSB), respectively. The Y216R/R398A mutation displayed elevated MSC transamination activity compared to wild-type, while mutations such as W18H, E27G, G36S, and H279F showed comparable MSC transamination activity to wild-type hKYAT1 (**Figure 4A**). Substitution of the aromatic residue at position 279 with a basic amino acid (H279F) resulted in a two-fold increase in MSC transamination activity in recombinant proteins as compared to wild-type (**Figure 4B**). This increased transamination activity in the H279F mutant was reflected in the MSC kinetics, with K_m_ and K_cat_ values of 15.7+/−5.4 and 0.8+/−0.15 mM, and 843 and 343 s^−1^ for wild-type and H279F, respectively (**Figure 4C, Table 1**). In contrast, none of the tested hKYAT1-mutants showed any significant transamination activity of SeMet, compared to wild-type hKYAT1, although hKYAT1-overexpressed cell lysates (wild-type or mutant) demonstrated significant transamination activity as compared to control cell lysates (**Figure 4D**). These results suggest that hKYAT1 preferentially transaminates MSC over SeMet (**Figure 4E**). The SeMet kinetics for wild-type and mutant KYAT1 further support this preference, showing K_m_ and K_cat_ values of 17.9+/−6.8 and 119.6+/−213 mM, and 175 and 837 s^−1^ for wild-type hKYAT1 and H279F, respectively (**Figure 4F, Table 1**).

**Figure 4.**
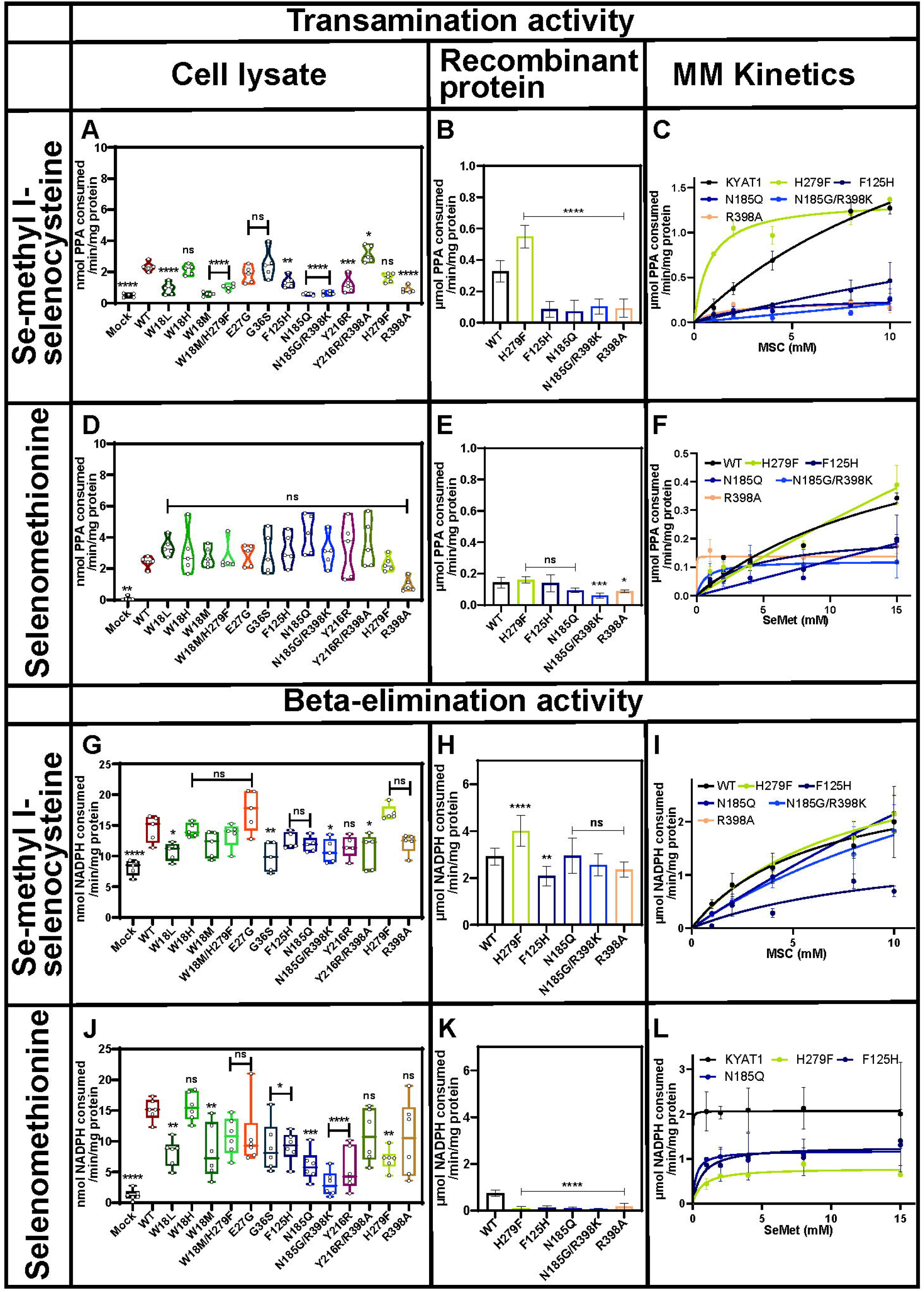
Mutation of hKYAT1 at the H279 substrate stabilization site alters transamination and β-elimination activities with MSC but not SeMet. The transamination efficacy of thirteen mutated hKYAT1 variants was assessed for Se-conjugated compounds, MSC (5 mM) and SeMet (5 mM). Panels **A** and **D** show activity with cell lysates containing 20 µg of crude protein, while panels **B** and **E** present data for five recombinant mutant proteins. Panels **C** and **F** show Michaelis-Menten kinetics with increasing MSC and SeMet for wild-type and five-mutants, using 200 ng of recombinant protein. β-elimination activity was tested in panels **G** and **J** with cell lysates (20 µg of crude protein), in panels **H** and **K** with recombinant mutant proteins (200 ng), and in panels **I** and **L** with Michaelis-Menten kinetics for MSC and SeMet. Unstable β-elimination activity was observed in the N185G/R398K mutant with MSC, while no detectable activity was seen with SeMet as a substrate in both N185G/R398K and R398A mutants. Statistical significance was calculated using one-way ANOVA with a 95% confidence interval, followed by Dunnett’s multiple comparison test (mean ± s.d., *P<0.05, n ≥5 per group).

In addition to cystine S-conjugates, KYAT1 also metabolizes Se-conjugates including MSC and SeMet, exhibiting β-elimination activity that cleaves MSC and SeMet into MS. This is attributed to the weaker C-Se bond compared to the C-S bond. Our data demonstrated that hKYAT1 induction enhanced β-elimination activity two-fold with MSC. Mutations such as W18L, G36S, N185G/R398K, and Y216R/R398A decreased β-elimination activity as compared to wild-type in the cell lysates (**Figure 4G**), while all other mutations showed similar β-elimination activity to wild-type in cell lysates (**Figure 4G**). In recombinant proteins, the H279F mutant exhibited a significant increase in β-elimination activity compared to wild-type (**Figure 4H**). This elevated β-elimination activity for both wild-type and the H279F mutant in recombinant proteins was reflected in their K_m_ and K_cat_ values (**Figure 4I, Table 1**).

SeMet was also tested for β-elimination activity in both wild-type and mutant hKYAT1. We observed elevated β-elimination activity for SeMet in hKYAT1-overexpressing cell lysates compared to control lysates. Different hKYAT1 mutants exhibited varying abilities to cleave SeMet into MS via β-elimination activity (**Figure 4J**). Results obtained with recombinant mutant and wild-type enzymes indicated that SeMet is not a preferred β-elimination substrate for hKYAT1 compared to MSC (**Figure 4K)**, This preference was further confirmed by the K_m_, K_cat,_ and K_cat_/K_m_ values for SeMet (**Figure 4L, Table 1**). A comparison of β-elimination activity between HEPG2 cell lysates and recombinant proteins for MSC and SeMet confirmed that MSC is favored as a substrate of hKYAT1 over SeMet.

## Discussion

KYAT1 plays a crucial role in recovering α-keto acid analogs from several essential amino acids through transamination reactions (9, 27). It is a key enzyme in the metabolism of the essential amino acid tryptophan, serving as the primary source for *de novo* nicotinamide adenine dinucleotide (NAD) biosynthesis (17). It transfers an amino group from kynurenine (an amino group donor in the Trp pathway) to α-ketoglutarate (an amino group acceptor), producing anthranilic acid and glutamate. The activity of hKYAT1, which exhibits broad substrate specificity, is regulated by multiple factors, including substrate availability, cofactors, and cellular energy status. As such, this crucial enzyme is involved in a range of physiological and pathological processes, including neurodegeneration, inflammation, and cancer.

The presence of aromatic residues in the ligand-binding region of hKYAT1 is crucial for substrate recognition (23, 25). Mutation analysis of catalytic, ligand-binding, and substrate-stabilization sites is therefore essential to understand the enzyme’s specificity toward its broad range of substrates. In this study, we identified and mutated several key residues in hKYAT1 and systematically characterized the effects of each substitution on a large array of amino acid substrates. It is well established that KYAT1 prefers different α-keto acids (amino group acceptors) depending on the substrate (9). For example, KYAT1 utilizes α-keto-γ-methylthiobutyric acid (KMB) and 2-keto-butyric acid (KBA) for the transamination of phenylalanine and kynurenine, respectively, while both α-ketoglutarate (αKG) and KMB are required for β-elimination activity with MSC. We carefully evaluated various amino group acceptors (KMB, KBA, αKG, pyruvate, phenylpyruvate (PPA), and glyoxylate) for their transamination activity with different substrates. Our results demonstrated that KMB, KBA, and PPA were the primary amino group acceptors for hKYAT1, with αKG showing the least activity.

hKYAT1 is also highly efficient at metabolizing α-keto acids into their corresponding amino acids, catalyzing reactions such as the conversion of phenylpyruvate to L-Phe, and indole-3-pyruvate (IPA) to L-Trp (3, 28). Previous studies have shown that hKYAT1 prefers transamination of IPA to L-Trp over transamination of L-Trp to IPA (3). These findings align with our results, which demonstrated increased transamination of aromatic amino acids like Phe and Trp. Aromatic amino acids are essential for various physiological functions, including quenching reactive oxygen species (ROS), providing neuroprotection, and facilitating signal transmission (11, 29, 30). Defects in their metabolism can have significant repercussions on human health (31).

In this study, we demonstrated that mutations in the substrate binding site (H279) and catalytic site (E27) significantly modulated the transamination activity of hKYAT1 for all aromatic amino acids. Rossi *et al* previously proposed that E27 plays a critical role in the catalytic site of KYAT1, where its mutation reduced enzyme activity and caused hypertension in rats (10, 26). However, we observed the opposite since E27 increased the activity. This could be due to greater flexibility in the α1-helix, potentially enhancing the adaptability of the active site for larger substrates (32). The E27G mutation increased enzymatic activity across most substrates, particularly Gln, but showed only moderate activity with Asp. These results emphasize the role of Gln-derived metabolites, such as αKG, in fueling the TCA cycle and maintaining anaplerosis (30, 33). Comparison of the crystal structure of hKYAT1 in complex with different substrates revealed that the lateral movement of the α1-helix is necessary for accommodating larger ligands, but not small ones (23). Residue H279 forms a hydrogen bond with bound substrates and plays a key role in stabilizing ligands, as demonstrated in the crystal structure of hKYAT1 (25). Our findings show that the H279F mutation significantly increased hKYAT1 activity toward Phe, Trp, His, and MSC, but reduced activity towards Tyr, Gln, Met, Gly, Cyss, and Kyn.

We observed in a few cases contradictory transamination activities between cell lysates and recombinant proteins. This variation could be attributed to the presence of other enzymes present in the whole-cell lysates, such as ω-amidase, tyrosine transaminase (TAT), and cytosolic malate dehydrogenase (MDH1), which are involved in the Trp salvage pathway (3). Additionally, since KYAT1 functions as a homodimer, we postulate that mutant KYAT1 subunits may dimerize with endogenous wild-type KYAT1, potentially affecting transamination activity in cell lysates. In recombinant proteins, the absence of such interference allowed for a clearer assessment of mutation effects, such as the increased activity of the H279F mutant with Trp, His, and MSC. Residue F125 is conserved across mammalian KYAT proteins, but in human KYAT2 and KYAT3, F125 is replaced by Tyr. In KYAT2, the Y125F mutation resulted in a twenty-fold reduction in β-lytic activity (34). Our findings indicate that the F125H mutation significantly affected substrate binding across all tested amino acid substrates. Residue W18, located at the hinge of the α1-helix, plays a critical role in ligand recognition and stabilization (23). Mutation at this residue inactivated hKYAT1 with most amino acids except Asp. Interestingly, the W18H mutation exhibited increased transamination activity for Kyn and Leu, although expression levels were double that of the wild-type (**Figure S1**). Residue G36, located at the entrance of the substrate channel, also plays a critical role, as demonstrated by the G36S mutation, which showed two-fold higher transamination activity for Leu compared to the wild-type. Mutations at N185 and R398, both key residues in the ligand-binding regions (23, 25), revealed additional insights. The N185Q/G mutants exhibited moderate activity for most substrates, with notably higher activity for L-Phe. In contrast, the R398A and Y216R mutations resulted in complete inactivation of hKYAT1, confirming the critical role of these residues in catalysis.

Transamination of kynurenine (Kyn) to kynurenic acid (Kyna) is a key step in the kynurenine pathway (KP), which plays an important role in inflammatory responses, immunoregulation, and psychiatric disorders (35). Kyna is considered neuroprotective, suppressing several inflammatory pathways and playing a significant role in immune responses by activating the aryl hydrocarbon receptor (AhR) (35). Interestingly, while the transamination of Kyn increased in hKYAT1 mutants in cell lysates, recombinant hKYAT1 mutants showed the opposite trend. This discrepancy suggests that Kyna production may not occur under recombinant conditions and could require additional factors, such as cofactors or other enzymes, to drive the reaction. It has been reported that Kyna does not cross the blood-brain barrier (BBB), whereas Kyn does, suggesting that Kyna is synthesized in the brain (36).

Our results demonstrated significantly higher transamination and β-elimination activity for the H279F mutants with MSC, but not with SeMet. The transamination and β-elimination of MSC result in the formation of β-methylselenopyruvate (MSP) and methylselenol (MS), respectively (36). In contrast, the transamination and β-elimination of SeMet generate α-keto-γ-methylselenobutyrate (KMSB) and methylselenol (MS), respectively (37). Both MSC and SeMet metabolites are known to modulate tumor cell growth and possess chemopreventive activities (37, 38). Specifically, MSP has been reported to inhibit histone deacetylases (HDACs), as it structurally resembles butyrate, a potent HDAC inhibitor (37). Additionally, the β-elimination products of MSC and SeMet can alter the redox balance, leading to changes in cell signaling pathways and the induction of proapoptotic genes in various cancers (39). Zeng *et al* (2006) reported that MS downregulates the BCL2-related protein A1 (BCL2A1), leading to increased cytochrome C release into the cytoplasm and initiation of apoptotic events (40, 41). Moreover, MSC has been shown to induce apoptosis by activating multiple kinase signaling pathways and by participating in the regulation of the NF-κB signaling pathway during inflammatory responses (42). In our study, elevated β-elimination activity with MSC was observed in both the E27G and H279F mutants in cell lysates. Furthermore, the recombinant H279F mutant exhibited higher β-elimination activity with MSC compared to the wild-type. In contrast, the other recombinant proteins tested displayed β-elimination activity similar to the wild-type, with the exception of F125H. Interestingly, only the wild-type enzyme showed moderate β-elimination activity with SeMet, further confirming that KYAT1 preferentially utilizes MSC as a substrate over SeMet, consistent with previous findings (43).

Despite transfecting equal quantities of mutant KYAT1 plasmids into HEPG2 cells, we observed variability in expression levels among the mutants (**Figures S1A, S1B**). Interestingly, the effect of these mutations appeared to be substrate-dependent. For example, while the H279F, R398A, and F125H mutants showed a two-fold increase in protein expression as compared to wild-type, this did not directly correlate with enzyme activity. Specifically, the H279F mutant displayed higher transamination activity toward Phe, whereas the R398A and F125H mutants showed no detectable transamination activity toward Phe. Additionally, despite its expression being twice as high as wild-type, the R398A mutant showed no significant activity toward several amino acid substrates (**Figures S1A, S1B**). These observations were further validated in recombinant hKYAT1 assays, where an equal quantity of KYAT1 mutants was used in all transamination and β-elimination assays. This underscores the importance of substrate specificity and the differential impact of mutations on hKYAT1 enzyme activity.

In conclusion, our study demonstrates that the transamination and β-elimination activity of hKYAT1 mutants varies depending on the substrate and the residues involved. We identified two key mutants, H279F and E27G, that significantly enhanced transamination activity toward several KYAT1-preferred substrates, including MSC. The ability to modulate transamination activity via mutation in vitro suggests that modification of hKYAT1 properties pharmacologically or genetically can be leveraged therapeutically in vivo so to enhance antineoplastic metabolism.

## Materials and Methods

### Chemicals and reagents

L-Tryptophan (L-Trp), L-Kynurenine (L-Kyn), L-Leucine (L-Leu), L-Alanine (L-Ala), L-Glutamine (L-Gln), L-Cystine (L-Cyss), L-Histidine (L-His), L-Asparagine (L-Asn), L-Aspartic acid (L-Asp), Phenylpyruvic acid (PPA), α-Keto-γ-methylthiobutyric acid sodium salt (KMB), 2-keto-butyric acid (KBA), Se-Methylselenocysteine hydrochloride (MSC), Dimethyl-2-oxoglutarate (α-KG), Glycine (Gly), Proline (Pro), L-Phenyl alanine (L-Phe), dL-Methionine (dL-Met), dL-Tyrosine (dL-Tyr), 2-Amino-2-methyl-1,3-propanediol, Pyridoxal 5’-phosphate hydrate (PLP), Phenylmethanesulfonyl fluoride (PMSF), RIPA buffer, Protease inhibitor cocktail mix, N-N-Dimethyl formamide, Potassium dihydrogen phosphate, Di-potassium hydrogen phosphate, EDTA, Sodium arsenate dibasic heptahydrate and Sodium hydroxide were purchased from Sigma-Aldrich (St. Louis, MO, USA). L-Selenomethionine was purchased from Santa Cruz (Dallas, TX, USA). NADPH was purchased from Acros Organics (Geel, Belgium). Lipofectamine 3000 was purchased from Invitrogen (Camarillo, CA, USA). Page Ruler Plus Prestained protein ladder was purchased from Thermo Fischer Scientific (Rockford, IL, USA). Plasmid pEGFP-N1 (Clontech, Takara Bio Inc, Mountain View, CA, USA) was kindly provided by Dr. Gildert Lauter and Dr. Peter Swoboda, Department of Biosciences and Nutrition, Karolinska Institute. Mammalian TrxR1 was purchased from Sigma-Aldrich (Darmstadt, Germany).

### Cell culture and growth conditions

HEPG2 cells were purchased from ATCC (Wesel, Germany) and maintained in EMEM (ATCC) supplemented with 10% heat-inactivated fetal bovine serum (FBS; Gibco, Paisley, UK) under 5% CO_2_ at 37°C without antibiotics. Cells were regularly tested for mycoplasma contamination using MycoAlert (Lonza, Boston, MA, USA). Cell counting was performed using a TC 20™ automated cell counter (Bio-Rad, Portland, ME, USA).

### Cloning and cellular overexpression of hKYAT1 and mutant variants

The cDNA encoding full-length hKYAT1 or its mutated variants was cloned into the mammalian expression vector pEGFP-N1 (Clontech, Takara Bio Inc., CA, USA). The human PGK promoter and puromycin resistance gene sequences were amplified from pLKO.1 (Sigma Aldrich, Darmstadt, Germany) and inserted into the pEGFP-N1 backbone containing the hKYAT1 expression vector. All mutants were confirmed by Sanger sequencing. An empty vector was created by inserting the PCR-amplified puromycin resistance gene expression cassette into pEGFP-N1. Protein sequences used in this study are listed in **Table S2**.

### Cloning, expression, and purification of recombinant hKYAT1 and mutated variants

The production and isolation of recombinant proteins were carried out using full-length hKYAT1 cDNA, which was amplified and cloned into the pET-23a expression vector (Novagen, Cambridge, UK) using the ligation-independent “Fast Cloning” method (44). All mutants were confirmed by Sanger sequencing. The plasmids were transformed into T7 Express Competent E. coli (New England Biolabs, USA), and overnight cultures were grown in Luria-Bertani broth (LB) medium containing 100 mg/ml ampicillin at 37°C, shaking at 120 rpm, and until OD600 reached 0.8. After cooling the cultures for 30 minutes at 4°C, protein expression was induced with 0.5 mM IPTG and grown overnight at 16°C. Harvested cells were stored at −20°C until use. All protein purification steps were performed at 4°C. Cell pellets were lysed in PBS buffer and subjected to ultrasonication. After centrifugation at 60,000 g for 30 min, protein in the supernatant was affinity purified using a HisTrap™ HP column (Cytiva, Uppsala, Sweden). Proteins were concentrated, and size exclusion chromatography was performed using a Superdex 200 increase 10/300 column (Cytiva, Uppsala, Sweden). Purified proteins were flash frozen in liquid nitrogen, and stored at −80°C.

### Overexpression of wild-type and mutated hKYAT1 variants in HEPG2 cells

HEPG2 cells were seeded at 400 cells/mm^2^ in 6-well plates. Twenty hours later, cells were transfected with 2 µg of plasmids DNA encoding wild-type or mutant hKYAT1 using Lipofectamine 3000 (Invitrogen, USA). An empty vector served as a negative control. Cells were harvested 48 hours post-transfection, and protein expression was verified by Western blot. Whole-cell lysates were used for subsequent enzyme assays.

### Determination of protein concentration

Samples were lysed on ice for 30 min in RIPA buffer containing 1 mM PMSF and 1% protease inhibitor cocktail mix, followed by sonication at 4°C for 30 seconds. Lysates were centrifuged at 13,000 rpm for 10 min at 4°C, and supernatants were collected. Protein concentration was determined using the Pierce™ Bicinchoninic acid (BCA) Protein Assay Kit (Thermo Fischer Scientific, Rockford, IL, USA) following the manufacturer’s instructions.

**Transamination assay for L-Phe** was performed according to a previously published protocol (36).

**Transamination assays for L-Leu, L-Ala, L-Cyss, L-Gln, dL-Met, Gly, Pro, SM, and MSC** were performed according to a previously published protocol (45).

**Transamination of L-Trp, dL-Tyr, and L-His** was measured following a modified version of the published protocol (46). Briefly, 20μg of whole-cell lysate or 200 ng of purified recombinant hKYAT1 was added to 100 µL of reaction mixture containing 3 mM amino acid, 5 mM KMB, 70 µM PLP, and 300 mM ammediol-HCl buffer (pH 9.6). After incubation at 37°C for 30 min, the reaction was terminated by adding 7 µL of 50% trichloroacetic acid (TCA). The mixture was thereafter vortexed and centrifuged for 2 min at maximum speed. Then, 250 µl of 1 M arsenate-borate reagent (pH 6.0) was added to 50 µl of the reaction mixture, and the reaction mixture was vortexed and incubated for an additional 30 min at room temperature. Absorbance was acquired at 292 nm for L-His (imidazolepyruvate, ε=11,300), 310nm for dL-Tyr (p-hydroxyphenylpyruvate, ε=10,700), and 330nm for L-Trp (indolepyruvate, ε=10,800) in a spectrophotometer (PowerWave HT, BioTek, USA). A mixture in which KMB was added immediately before adding 50% TCA was used as a blank.

**Transamination of L-Kyn to L-Kyna** with KBA as α-keto acid was performed using a 50 µL reaction mixture that contained 3 mM L-Kyn, 5 mM KBA, 200 µM PLP, and 300 mM ammediol-HCl buffer (pH 9.6), to which 40 µg of whole-cell lysate or 400 ng of recombinant hKYAT1 was added. The assay mixture was incubated at 37°C for six hours, and the reaction was terminated by adding 3.5 µL of 50% TCA. The assay mixture was thereafter vortexed and centrifuged for 2 min at maximal speed. Then, 180 µl 500 mM phosphate buffer pH 7.5 was added to 20 µl of the reaction mixture, and absorbance was read immediately at 330 nm in a spectrophotometer (PowerWave HT, BioTek, USA). Under this condition, the molar extinction coefficient is 8,850 M^−1^ cm^−1^ for L-Kyna (47). A mixture in which KBA was added immediately before adding 50% TCA was used as a blank.

**Beta-elimination activity was measured for MSC and SM** according to a previously published method (36). In all assays, 20 μg of whole-cell lysates or 200 ng of recombinant proteins was used unless otherwise specified.

### Immunoblotting

Western blot analysis was performed by loading 20 µg protein isolated from whole-cell lysates onto a 4-20% Mini-PROTEAN gel, followed by transfer to a PVDF membrane (Bio-Rad, USA). Membranes were blocked with 5% milk in TBST for two hours, then incubated with primary antibodies against KYAT1 (PA5-51313, 1:1000 in 5% BSA) (Sigma, Germany) overnight at 4°C. Vinculin (Millipore, USA) was used as a loading control. Membranes were washed and incubated for one hour with a secondary antibody, polyclonal swine anti-rabbit immunoglobulins/HRP (P0399) diluted 1:10,000 in 5% milk (DAKO, Denmark). Membranes were subsequently washed three times with TBST, and blot images were acquired using the Odyssey Fc Imaging System (LI-COR, USA). Protein levels were quantified using Odyssey Image software (LI-COR Biosciences, USA), by normalizing primary antibody signals to vinculin.

### Statistical analysis

Results are expressed as mean ± SD (n ≥ 3) and are displayed using box and whisker plots or violin plots. Statistical analyses were conducted using one-way ANOVA with a 95% confidential interval, followed by Dunnett’s multiple comparison test. Statistical significance was set at p < 0.05. Data were analyzed using GraphPad Prism (version 10.1.2, GraphPad Software Inc., USA).

## Data Availability Statement

Data are available from the corresponding author upon request.

## Conflicts of Interest

Mikael Björnstedt is listed as an inventor in a patent application for *i*.*v*. Use of inorganic selenium in cancer patients and hold shares in SELEQ OY, a company involved in the development of Se-based formulations for prevention and treatment.

## Acknowledgments

This study was supported by grants from Swedish Cancer Society [23 2796 Pj 01 H], Radium Hemmets Research Funds [231082], Center for Innovative Medicine [FoUI-976014] to M.B, and by grants from Swedish Cancer Society [21 1605 Pj 01 H and 24 3775 Pj 01 H], Cancer and Allergy Foundation [10399], Swedish Research Council [2021-05061 and 2018-02874], and the King Gustaf V Jubileum Foundation [244092] to A.A.

## Supporting information captions

**Table S1: Transamination and β-elimination activity of wild-type and mutant hKYAT1 with various amino acid substrates in crude cell extracts**

**Table S2: Protein sequences of KYAT wild-type and mutants.**

**Figure S1. Residual changes in hKYAT1 lead to varying protein expression in HEPG2 cells.** Western blot analysis and quantification of wild-type and 13 different mutants. An equal amount (20 μg) of whole-cell lysate was loaded onto the gel. Statistical significance was determined using one-way ANOVA with a 95% confidence interval, followed by Dunnett’s multiple comparison test (mean ± s.d., *P<0.05, n=3 per group).

**Figure S2. Mutation of hKYAT1 does not alter transamination activity with different amino acids.** Transamination efficacy of thirteen different mutated variants of hKYAT1 was assessed for Tyr (3 mM), Asp (3 mM), Cyss (3 mM) Pro (3 mM), Ala (3 mM), & Gly (3 mM). **(A, C, E, G, I & K)** Cell lysates containing 20 μg of crude protein, and **(B, D, F, H, J & L)** wild-type and five recombinant mutant proteins, each at 200 ng, were used in the assays. Statistical significance was determined using one-way ANOVA with a 95% confidence interval, followed by Dunnett’s multiple comparisons test (mean ± s.d., *P<0.05, n ≥5 per group).

## Notes

### Competing Interest Statement

Mikael Bjornstedt is listed as an inventor in a patent application for i.v. Use of inorganic selenium in cancer patients and hold shares in SELEQ OY, a company involved in the development of Se-based formulations for prevention and treatment.

