## Supporting Information for "Mutational insights into human kynurenine aminotransferase 1: modulation of transamination and β-elimination activities across diverse substrates"

**Table S1: Transamination and β-elimination activity of wild-type and mutant hKYAT1 with various amino acid substrates in crude cell extracts**

| **Transamination activity in crude cell extracts** | | | | | | | | | | | | | | | | |
|---|---|---|---|---|---|---|---|---|---|---|---|---|---|---|---|---|
| | **L-Phe** | **L-Trp** | **L-Kyn** | **L-Gln** | **L-Asn** | **L-His** | **dL-Met** | **L-Leu** | **MSC** | **SeMet** | **dL-Tyr** | **L-Asp** | **L-Cyss** | **L-Ala** | **L-Pro** | **Gly** |
| **Mock (empty plasmid)** | 1.2+/-0.3 | 0.3+/-0.2 | 2.0+/-0.7 | 1.1+/-0.4 | 0.03+/-.02 | 0.6+/-0.2 | 1.1+/-0.4 | 0.8+/-0.3 | 0.6+/-0.2 | 0.1+/-0.1 | 0.6+/-0.3 | 0.3+/-0.02 | 0.4+/-0.1 | 0.3+/-0.1 | 1.5+/-0.4 | 0.4+/-0.2 |
| **Wild-type** | 11.2+/-2.7 | 2.1+/-0.9 | 5.1+/-1.6 | 8.7+/-1.7 | 1.0+/-0.3 | 4.8+/-1.2 | 10.8+/-1.4 | 4.9+/-2.1 | 2.3+/-0.3 | 2.4+/-0.3 | 4.0+/-1.7 | 0.8+/-0.1 | 2.1+/-0.6 | 2.4+/-0.8 | 3.3+/-0.8 | 2.6+/-0.4 |
| **W18L** | 2.1+/-0.4 | 0.4+/-0.3 | 8.9+/-3.9 | 2.2+/-0.6 | 2.0+/-0.4 | 0.2+/-0.1 | 7.4+/-2.9 | 0.9+/-0.9 | 0.9+/-0.4 | 3.4+/-0.5 | 0.6+/-0.2 | 2.0+/-0.33 | 0.8+/-0.3 | 0.4+/-0.1 | 1.6+/-0.5 | 2.4+/-0.4 |
| **W18H** | 7.0+/-1.5 | 1.4+/-0.6 | 13.2+/-2.4 | 4.3+/-1.2 | 1.7+/-0.3 | 1.1+/-0.2 | 2.7+/-1.1 | 7.4+/-2.3 | 2.1+/-0.3 | 3.1+/-1.3 | 1.1+/-0.2 | 2.1+/-0.5 | 2.5+/-1.1 | 1.2+/-0.8 | 1.1+/-0.5 | 0.6+/-0.3 |
| **W18M** | 4.4+/-0.6 | 0.4+/-0.2 | 4.9+/-2.2 | 0.6+/-0.1 | 1.1+/-0.5 | 1.2+/-0.3 | 2.3+/-1.3 | 1.8+/-0.4 | 0.6+/-0.1 | 2.8+/-0.6 | 0.4+/-0.04 | 2.0+/-0.5 | 2.2+/-0.7 | 4.0+/-1.0 | 2.8+/-0.5 | 2.4+/-0.9 |
| **W18M/H279F** | 1.6+/-0.9 | 0.7+/-0.2 | 12.1+/-2.3 | 6.2+/-2.7 | 1.8+/-0.5 | 1.0+/-0.2 | 2.3+/-1.0 | 2.3+/-0.8 | 1.0+/-0.2 | 2.7+/-0.8 | 1.3+/-0.6 | 2.4+/-0.1 | 1.4+/-0.3 | 1.2+/-0.7 | 0.5+/-0.2 | 1.6+/-0.5 |
| **E27G** | 19.2+/-1.9 | 3.4+/-1.4 | 12.8+/-5.9 | 13.5+/-4.5 | 1.5+/-0.3 | 3.4+/-1.7 | 6.6+/-1.1 | 4.6+/-1.1 | 1.8+/-0.5 | 2.9+/-0.6 | 2.2+/-1.0 | 2.0+/-0.2 | 2.2+/-0.7 | 2.3+/-1.2 | 1.9+/-0.5 | 0.9+/-0.2 |
| **G36S** | 2.0+/-0.8 | 0.1+/-0.02 | ND | 5.3+/-3.3 | 1.2+/-0.3 | 1.3+/-0.4 | 2.2+/-0.3 | 13.6+/-2.8 | 2.5+/-0.9 | 3.0+/-1.2 | 0.5+/-0.2 | 1.9+/-0.5 | 3.0+/-1.2 | 2.6+/-0.8 | 3.8+/-1.3 | 3.2+/-1.2 |
| **F125H** | 1.4+/-0.2 | 0.4+/-0.2 | 14.4+/-1.9 | 3.2+/-0.6 | 1.6+/-0.2 | 0.9+/-0.2 | 7.9+/-3.6 | 1.5+/-0.5 | 1.4+/-0.4 | 3.2+/-0.9 | 1.2+/-0.3 | 2.2+/-0.5 | 2.2+/-0.7 | 2.7+/-1.1 | 2.8+/-0.7 | 2.1+/-0.8 |
| **N185Q** | 20.7+/-1.6 | 10.5+/-4.2 | 4.6+/-1.1 | 9.8+/-3.5 | 2.6+/-1.1 | 2.7+/-1.1 | 4.3+/-2.2 | 4.6+/-3.7 | 0.6+/-0.1 | 4.2+/-1.1 | 0.8+/-0.4 | 2.5+/-0.6 | 2.0+/-0.3 | 4.1+/-1.9 | 2.0+/-1.5 | 2.6+/-0.6 |
| **N185G/R398K** | 1.1+/-0.6 | 0.2+/-0.2 | 0.3+/-0.2 | 5.6+/-2.3 | 1.1+/-0.4 | 1.9+/-0.4 | 3.6+/-1.1 | 2.8+/-2.0 | 0.6+/-0.1 | 3.1+/-0.9 | 0.4+/-0.2 | 2.3+/-0.3 | 1.7+/-0.3 | 4.8+/-2.2 | 2.2+/-1.2 | 2.3+/-1.2 |
| **Y216R** | 1.0+/-0.2 | 0.2+/-0.1 | 6.5+/-3.0 | 4.5+/-2.2 | 0.3+/-0.1 | 0.1+/-0.1 | 4.1+/-1.6 | 5.1+/-1.6 | 1.2+/-0.5 | 3.3+/-1.6 | 0.2+/-0.1 | 0.7+/-0.2 | 2.9+/-1.3 | 2.3+/-0.8 | 3.0+/-1.8 | 2.2+/-1.2 |
| **Y216R/R398A** | 0.9+/-0.1 | 0.3+/-0.3 | 3.3+/-0.8 | 3.4+/-1.6 | 0.6+/-0.3 | 0.6+/-0.2 | 1.5+/-0.2 | 1.5+/-1.3 | 3.2+/-0.5 | 3.6+/-1.4 | 0.7+/-0.3 | 2.0+/-0.5 | 2.0+/-0.7 | 1.8+/-0.9 | 2.5+/-0.3 | 2.1+/-0.6 |
| **H279F** | 18.8+/-4.6 | 0.8+/-0.2 | 16.6+/-3.0 | 6.3+/-1.2 | 1.6+/-0.6 | 1.6+/-0.5 | 4.2+/-1.7 | 3.6+/-2.3 | 1.6+/-0.3 | 2.3+/-0.4 | 0.4+/-0.2 | 1.9+/-0.6 | 2.4+/-0.9 | 3.1+/-1.2 | 2.3+/-0.7 | 0.6+/-0.3 |

| | | | | | | | | | | | | | | | |
|---|---|---|---|---|---|---|---|---|---|---|---|---|---|---|---|
| **R398A** | 0.8+/-0.1 | 0.1+/-0.1 | 17.1+/-1.9 | 0.1+/-0.7 | 1.0+/-0.5 | 1.2+/-0.2 | 6.9+/-2.7 | 2.6+/-0.9 | 0.9+/-0.3 | 1.0+/-0.4 | 0.3+/-0.2 | 2.0+/-0.3 | 1.4+/-0.5 | 1.3+/-1.1 | 1.3+/-0.6 | 0.6+/-0.6 |

| **β-elimination activity in crude cell extracts** | | | | | | | | | | | | | | | |
|---|---|---|---|---|---|---|---|---|---|---|---|---|---|---|---|
| | **Mock** | **Wild type** | **W18L** | **W18H** | **W18 M** | **W18M/ H279F** | **E27G** | **G36S** | **F125H** | **N185Q** | **N185G/ R398K** | **Y216R** | **Y216R/ R398A** | **H279F** | **R398A** |
| **MSC** | 8.1+/-1.5 | 14.5+/-1.5 | 10.7+/-1.3 | 14.2+/-1.0 | 11.9+/-1.9 | 13.7+/-2.1 | 17.5+/-3.0 | 9.9+/-2.2 | 11.3+/-1.7 | 10.9+/-2.3 | 10.1+/-2.2 | 11.5+/-1.6 | 8.7+/-3.0 | 16.5+/-1.3 | 12.1+/-1.5 |
| **SeMet** | 1.3+/-0.8 | 15.2+/-1.6 | 8.2+/-2.0 | 15.6+/-2.1 | 8.4+/-4.0 | 10.8+/-2.8 | 10.8+/-4.7 | 9.1+/-3.7 | 9.1+/-2.2 | 6.1+/-2.2 | 3.1+/-1.8 | 5.5+/-3.1 | 10.9+/-3.9 | 7.1+/-1.6 | 10.5+/-5.5 |

**Table S2: Protein sequences of KYAT wild-type and mutants.**

| | **Protein sequences** |
|---|---|
| **Wild-type** | MAKQLQARRLDGIDYNPWVEFVKLASEHDVVNLGQGFPDFPPPDFAVEAFQHAVSGDFMLNQYTKTFGYPPLTKILASFFGELLGQEIDPL[illegible]<br>VFSREELELVASLCQQHDVVCITDEVYQWMVYDGHQHISIASLPGMWERTLTIGSAGKTFSATGWKVGWVLGPDHIMKHLRTVHQNSVFH<br>AIPVSIFYSVPHQKHFDHYIRFCFVKDEATLQAMDEKLRKWKVEL |
| **W18L** | MAKQLQARRLDGIDYNP**L**VEFVKLASEHDVVNLGQGFPDFPPPDFAVEAFQHAVSGDFMLNQYTKTFGYPPLTKILASFFGELLGQEIDPLR<br>VFSREELELVASLCQQHDVVCITDEVYQWMVYDGHQHISIASLPGMWERTLTIGSAGKTFSATGWKVGWVLGPDHIMKHLRTVHQNSVFH<br>AIPVSIFYSVPHQKHFDHYIRFCFVKDEATLQAMDEKLRKWKVEL |
| **W18H** | MAKQLQARRLDGIDYNP**H**VEFVKLASEHDVVNLGQGFPDFPPPDFAVEAFQHAVSGDFMLNQYTKTFGYPPLTKILASFFGELLGQEIDPLR<br>VFSREELELVASLCQQHDVVCITDEVYQWMVYDGHQHISIASLPGMWERTLTIGSAGKTFSATGWKVGWVLGPDHIMKHLRTVHQNSVFH<br>AIPVSIFYSVPHQKHFDHYIRFCFVKDEATLQAMDEKLRKWKVEL |
| **W18M** | MAKQLQARRLDGIDYNP**M**VEFVKLASEHDVVNLGQGFPDFPPPDFAVEAFQHAVSGDFMLNQYTKTFGYPPLTKILASFFGELLGQEIDPL[illegible]<br>VFSREELELVASLCQQHDVVCITDEVYQWMVYDGHQHISIASLPGMWERTLTIGSAGKTFSATGWKVGWVLGPDHIMKHLRTVHQNSVFH<br>AIPVSIFYSVPHQKHFDHYIRFCFVKDEATLQAMDEKLRKWKVEL |
| **W18M/H279F** | MAKQLQARRLDGIDYNP**M**VEFVKLASEHDVVNLGQGFPDFPPPDFAVEAFQHAVSGDFMLNQYTKTFGYPPLTKILASFFGELLGQEIDPL[illegible]<br>VFSREELELVASLCQQHDVVCITDEVYQWMVYDGHQHISIASLPGMWERTLTIGSAGKTFSATGWKVGWVLGPDHIMKHLRTVHQNSVF**F**<br>IPVSIFYSVPHQKHFDHYIRFCFVKDEATLQAMDEKLRKWKVEL |
| **E27G** | MAKQLQARRLDGIDYNPWVEFVKLAS**G**HDVVNLGQGFPDFPPPDFAVEAFQHAVSGDFMLNQYTKTFGYPPLTKILASFFGELLGQEIDPL[illegible]<br>VFSREELELVASLCQQHDVVCITDEVYQWMVYDGHQHISIASLPGMWERTLTIGSAGKTFSATGWKVGWVLGPDHIMKHLRTVHQNSVFH<br>AIPVSIFYSVPHQKHFDHYIRFCFVKDEATLQAMDEKLRKWKVEL |
| **G36S** | MAKQLQARRLDGIDYNPWVEFVKLASEHDVVNLGQ**S**FPDFPPPDFAVEAFQHAVSGDFMLNQYTKTFGYPPLTKILASFFGELLGQEIDPLR<br>VFSREELELVASLCQQHDVVCITDEVYQWMVYDGHQHISIASLPGMWERTLTIGSAGKTFSATGWKVGWVLGPDHIMKHLRTVHQNSVFH<br>AIPVSIFYSVPHQKHFDHYIRFCFVKDEATLQAMDEKLRKWKVEL |

| | |
|---|---|
| **F125H** | MAKQLQARRLDGIDYNPWVEFVKLASEHDVVNLGQGFPDFPPPDFAVEAFQHAVSGDFMLNQYTKTFGYPPLTKILASFFGELLGQEIDPL[illegible]<br>VFSREELELVASLCQQHDVVCITDEVYQWMVYDGHQHISIASLPGMWERTLTIGSAGKTFSATGWKVGWVLGPDHIMKHLRTVHQNSVFH<br>AIPVSIFYSVPHQKHFDHYIRFCFVKDEATLQAMDEKLRKWKVEL |
| **N185Q** | MAKQLQARRLDGIDYNPWVEFVKLASEHDVVNLGQGFPDFPPPDFAVEAFQHAVSGDFMLNQYTKTFGYPPLTKILASFFGELLGQEIDPL[illegible]<br>VFSREELELVASLCQQHDVVCITDEVYQWMVYDGHQHISIASLPGMWERTLTIGSAGKTFSATGWKVGWVLGPDHIMKHLRTVHQNSVFH<br>AIPVSIFYSVPHQKHFDHYIRFCFVKDEATLQAMDEKLRKWKVEL |
| **N185G/**<br>**R398K** | MAKQLQARRLDGIDYNPWVEFVKLASEHDVVNLGQGFPDFPPPDFAVEAFQHAVSGDFMLNQYTKTFGYPPLTKILASFFGELLGQEIDPL[illegible]<br>VFSREELELVASLCQQHDVVCITDEVYQWMVYDGHQHISIASLPGMWERTLTIGSAGKTFSATGWKVGWVLGPDHIMKHLRTVHQNSVFH<br>AIPVSIFYSVPHQKHFDHYI**K**FCFVKDEATLQAMDEKLRKWKVEL |
| **Y216R** | MAKQLQARRLDGIDYNPWVEFVKLASEHDVVNLGQGFPDFPPPDFAVEAFQHAVSGDFMLNQYTKTFGYPPLTKILASFFGELLGQEIDPL[illegible]<br>VFSREELELVASLCQQHDVVCITDEV**R**QWMVYDGHQHISIASLPGMWERTLTIGSAGKTFSATGWKVGWVLGPDHIMKHLRTVHQNSVFH<br>AIPVSIFYSVPHQKHFDHYIRFCFVKDEATLQAMDEKLRKWKVEL |
| **Y216R/**<br>**R398A** | MAKQLQARRLDGIDYNPWVEFVKLASEHDVVNLGQGFPDFPPPDFAVEAFQHAVSGDFMLNQYTKTFGYPPLTKILASFFGELLGQEIDPL[illegible]<br>VFSREELELVASLCQQHDVVCITDEV**R**QWMVYDGHQHISIASLPGMWERTLTIGSAGKTFSATGWKVGWVLGPDHIMKHLRTVHQNSVFH<br>AIPVSIFYSVPHQKHFDHYI**A**FCFVKDEATLQAMDEKLRKWKVEL |
| **H279F** | MAKQLQARRLDGIDYNPWVEFVKLASEHDVVNLGQGFPDFPPPDFAVEAFQHAVSGDFMLNQYTKTFGYPPLTKILASFFGELLGQEIDPL[illegible]<br>VFSREELELVASLCQQHDVVCITDEVYQWMVYDGHQHISIASLPGMWERTLTIGSAGKTFSATGWKVGWVLGPDHIMKHLRTVHQNSVF**F**<br>IPVSIFYSVPHQKHFDHYIRFCFVKDEATLQAMDEKLRKWKVEL |
| **R398A** | MAKQLQARRLDGIDYNPWVEFVKLASEHDVVNLGQGFPDFPPPDFAVEAFQHAVSGDFMLNQYTKTFGYPPLTKILASFFGELLGQEIDPL[illegible]<br>VFSREELELVASLCQQHDVVCITDEVYQWMVYDGHQHISIASLPGMWERTLTIGSAGKTFSATGWKVGWVLGPDHIMKHLRTVHQNSVFH<br>AIPVSIFYSVPHQKHFDHYI**A**FCFVKDEATLQAMDEKLRKWKVEL |

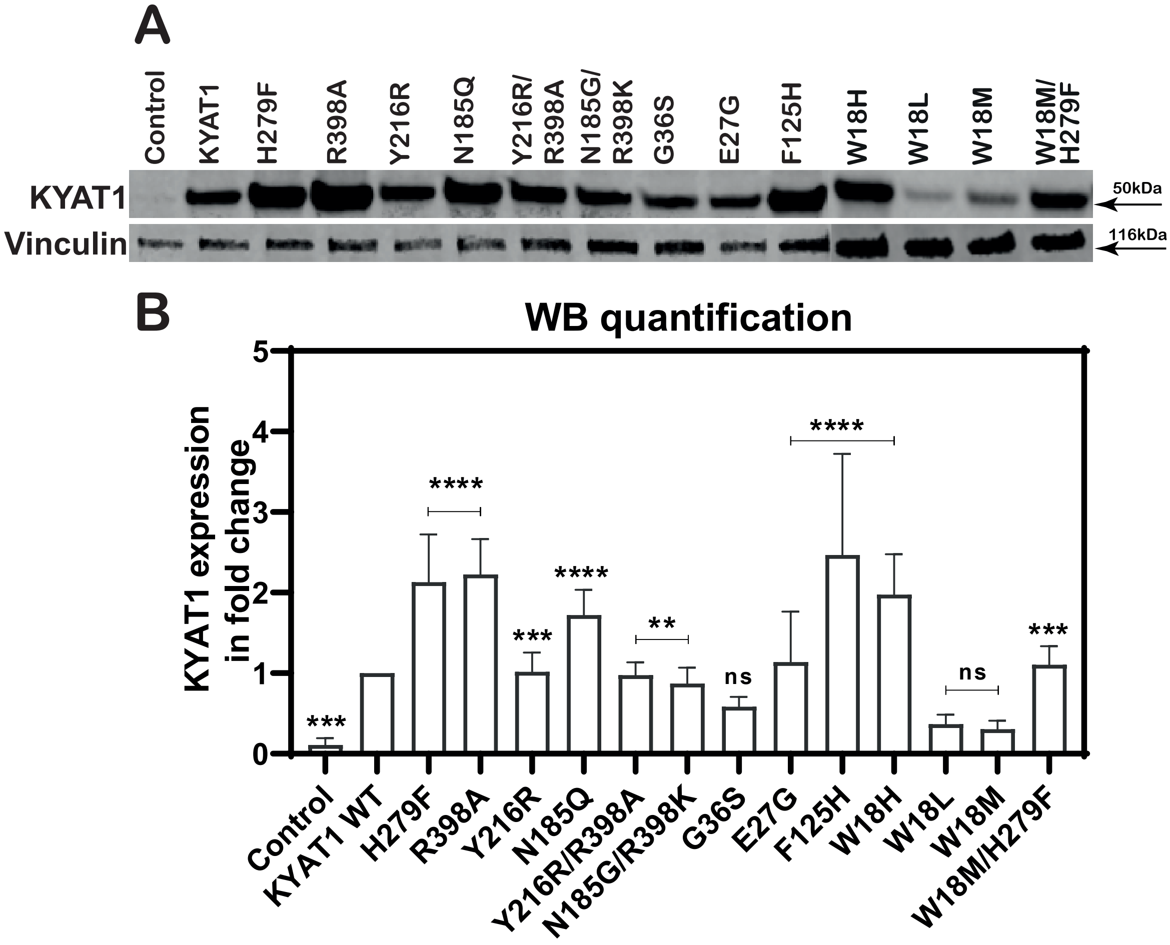


**Figure S1. Residual changes in hKYAT1 lead to varying protein expression in HEPG2 cells.**
**(A, B)** Western blot analysis and quantification of wild-type and 13 different mutants. An equal amount (20 µg) of whole-cell lysate was loaded onto the gel. Statistical significance was determined using one-way ANOVA with a 95% confidence interval, followed by Dunnett's multiple comparison test (mean ± s.d., *P<0.05, n=3 per group).

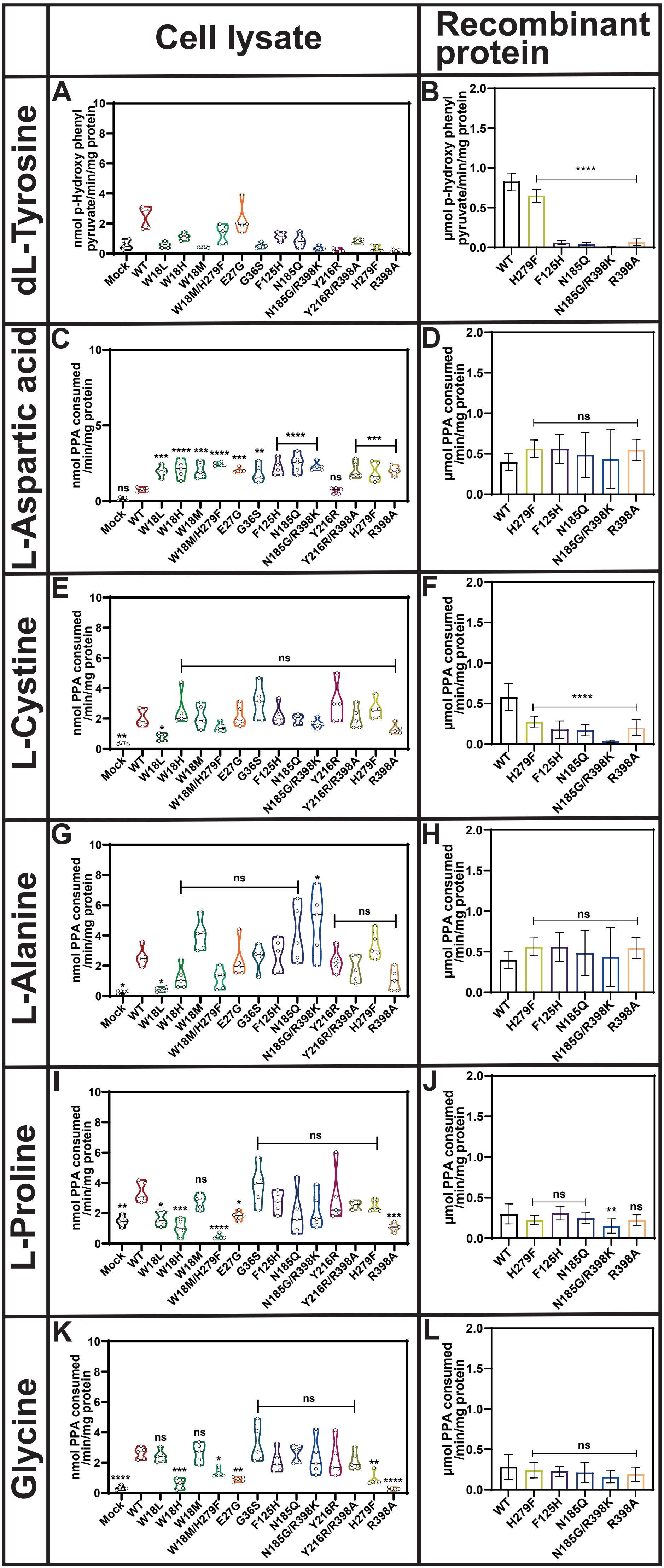

Cell lysate
Recombinant protein
dL-Tyrosine
L-Aspartic acid
L-Cystine
L-Alanine
L-Proline
Glycine
A
B
C
D
E
F
G
H
I
J
K
L
nmol p-Hydroxy phenyl pyruvate/min/mg protein
μmol p-Hydroxy phenyl pyruvate/min/mg protein
nmol PPA consumed /min/mg protein
μmol PPA consumed /min/mg protein
Mock
WT
W18L
W18H
W18M
W18M/H279F
E27G
G36S
F125H
N185Q
N185G/R398K
Y216R
Y216R/R398A
H279F
R398A
ns
*
**
***
****

**Figure S2. Mutation of hKYAT1 does not alter transamination activity with different amino acids.** Transamination efficacy of thirteen different mutated variants of hKYAT1 was assessed for Tyr (3 mM), Asp (3 mM), Cyss (3 mM) Pro (3 mM), Ala (3 mM), & Gly (3 mM). **(A, C, E, G, I & K)** Cell lysates containing 20 µg of crude protein, and **(B, D, F, H, J & L)** wild-type and five recombinant mutant proteins, each at 200 ng, were used in the assays. Statistical significance was determined using one-way ANOVA with a 95% confidence interval, followed by Dunnett's multiple comparisons test (mean ± s.d., *$P<0.05$, n ≥5 per group).
